# Bat influenza vectored NS1-truncated live vaccine protects pigs against heterologous virus challenge

**DOI:** 10.1101/2020.11.18.389254

**Authors:** Jinhwa Lee, Yonghai Li, Yuhao Li, A. Giselle Cino-Ozuna, Michael Duff, Yuekun Lang, Jingjiao Ma, Sunyoung Sunwoo, Juergen A. Richt, Wenjun Ma

## Abstract

Swine influenza is an important disease for the swine industry. Currently used whole inactivated virus (WIV) vaccines can induce vaccine-associated enhanced respiratory disease (VAERD) in pigs when the vaccine strains mismatch with the infected viruses. Live attenuated influenza virus vaccine (LAIV) is effective to protect pigs against homologous and heterologous swine influenza virus infections without inducing VAERD, but has safety concerns due to potential reassortment with circulating viruses. Herein, we used a chimeric bat influenza Bat09:mH3mN2 virus, which contains both surface HA and NA gene open reading frames of the A/swine/Texas/4199-2/1998 (H3N2) and six internal genes from the novel bat H17N10 virus, to develop modified live-attenuated viruses (MLVs) as vaccine candidates which cannot reassort with canonical influenza A viruses by co-infection. Two attenuated MLV vaccine candidates including the virus that expresses a truncated NS1 (Bat09:mH3mN2-NS1-128, MLV1) or expresses both a truncated NS1 and the swine IL-18 (Bat09:mH3mN2-NS1-128-IL-18, MLV2) were generated and evaluated in pigs against a heterologous H3N2 virus using the WIV vaccineb as a control. Compared to the WIV vaccine, both MLV vaccines were able to reduce lesions and virus replication in lungs and limit nasal virus shedding without VAERD, also induced significantly higher levels of mucosal IgA response in lungs and significantly increased numbers of antigen-specific IFN-γ secreting cells against the challenge virus. However, no significant difference was observed in efficacy between the MLV1 and MLV2. These results indicate that bat influenza vectored MLV vaccines can be used as a safe live vaccine to prevent swine influenza.

## Introduction

Influenza A viruses (IAVs) cause an acute respiratory disease ranging from mild pneumonia to death usually via co-infection with other respiratory pathogens in swine, leading to substantial economic losses to the swine industry each year. Currently, three major subtypes of IAVs including H1N1, H1N2 and H3N2 are endemic in US swine herds. However, there is extensive genetic and antigenic diversity of swine IAVs through antigenic drift, gene reassortment and introduction of IAVs from other hosts including human seasonal IAV into pigs [1–4]. Since the dynamic swine influenza viruses (SIVs) continue to evolve and generate novel IAVs, public health concerns increase with continual spillover of SIVs into human populations [5, 6]. The rapidly evolving diversity of IAVs hinders effective vaccine-mediated protection due to a deficiency of antigenic matching between vaccine strains and circulating viruses [7]. It is necessary and important to develop a new vaccine approach to achieve sufficient protection against circulating diverse SIVs and reducing the risk of zoonotic transmission to humans.

Currently, whole inactivated virus (WIV) vaccines are the most widely used vaccines with oil-in-water adjuvants in swine herds. The WIV vaccines are bivalent or multivalent products which contain combination of antigenically distinct H1 and H3 subtypes of viruses and usually require a two dose vaccination strategy delivered by intramuscular route. The adjuvanted WIV vaccines can provide sterilizing immunity by inducing robust neutralizing antibodies against antigenically similar HA strains [8, 9]. However, only partial protection is achieved by WIV vaccines against heterologous strains [9–11]. Moreover, vaccine associated enhanced respiratory disease (VAERD), characterized by absence of high avidity, cross-reactiv neutralizing antibodies as well as increased lung pathology, has been observed in pigs immunized with WIV vaccine followed by infection with antigenically distinct IAVs [11–15].

Live attenuated influenza virus vaccines (LAIVs) mimic a natural route of infection through intranasal administration. In contrast to WIV vaccines, LAIVs in swine provide improved cross-reactive immunity against antigenically distinct IAVs through inducing broader cell-mediated, humoral and mucosal immune responses without inducing VAERD [9, 16, 17]. Several approaches have been developed to attenuate wild type viruses to produce LAIV candidates, including elastase-dependent HA cleavage, truncated NS1 gene (ΔNS1) which is associated with suppression of type I interferon system, temperature-sensitive mutations in polymerase genes and codon-pair biased-deoptimization (CPBD) in HA and NA genes [18–22]. Among these approaches, a LAIV based on ΔNS1 was recently licensed as the first LAIV for swine in the US [23]. However, major concerns regarding the use of LAIVs are the reversion to a virulent phenotype of the vaccine strain over time by natural mutations or genome reassortment between vaccine strains and circulating field viruses.

Vectored vaccines are alternative approaches to produce IAV vaccines with potential application in multiple species. Replication-defective or replication-competent vectors infect cells and transport the recombinant genes and express the antigens of interest in infected cells, which allows the stimulation of cell mediated immunity. Furthermore, vectored vaccines can induce local immunity at the site of natural infection of IAVs through intranasal delivery [24, 25]. Human adenovirus type 5-vectored and alphavirus-like replicon particle vectored vaccines have been reported to provide complete protection against homologous IAV infection and partial protection against heterologous challenge in pigs [25–28].

Recently, two novel influenza A-like virus genome sequences were discovered from bat specimens and they were classified as H17N10 and H18N11 subtypes [29, 30]. Although the surface glycoproteins of these new bat influenza viruses are phylogenetically and structurally related to hemagglutinin (HA) and neuraminidase (NA) of conventional IAVs, they are functionally different from conventional HA and NA with lacking of hemagglutination and sialidase activities [30–32]. In addition, unrevealed cell tropism and lacking of suitable cultivation and replication systems of bat influenza viruses hindered its further characterization. Recently, we successfully rescued replicative chimeric bat influenza viruses which contain HA and NA coding regions from a conventional IAV with respective gene packaging signals of bat influenza viruses [33, 34]. Although the chimeric bat influenza viruses replicated to reasonable infectious titers in mammalian cells and mice, reassortment between chimeric bat influenza viruses and conventional IAVs was not observed in co-infection experiments [33–35]. This indicates that a chimeric bat influenza virus can be a good vaccine vector to overcome the potential risk of LAIV reassortment with conventional IAVs.

In the present study, we developed novel modified live-attenuated virus (MLV) vaccine candidates using a chimeric bat influenza virus as a vaccine vector containing a truncated NS1 gene. In addition, recombinant porcine IL-18 (rpIL-18) was introduced into this MLV as part of the MLV vaccine because IL-18 is known to induce strong cell mediated immunity (CMI) by stimulating T helper 1 (Th1) activation and IFN-γ induction [36–38], its effect on MLV vaccine efficacy was evaluated. The results showed that compared to WIV vaccine both MLV vaccines are able to reduce virus replication and pathology in lungs and limit virus nasal transmission without inducing VAERD after heterologous challenge in pigs. Furthermore, both MLV vaccines induced significant mucosal immunity and T-cell response against the challenge virus. However, a significant immunomodulatory effect of IL-18 after MLV2 vaccination was not observed.

## Materials and methods

### Ethical statement

Animal study (IACUC no.3668) was reviewed and approved by the Institutional Animal Care and Use Committee at Kansas State University and were performed in Biosafety Level 2+ animal facilities under guidance from the Comparative Medicine Group at Kansas State University.

### Viruses and vaccine preparation

The MLV with a truncated NS1 protein (Bat09:mH3mN2-NS1-128, MLV1) was generated via reverse genetics using six internal genes from the H17N10 A/little yellow-shouldered bat/Guatemala/164/2009 (Bat09) and two surface HA and NA gene coding regions from the H3N2 A/swine/Texas/4199-2/1998 (TX98) with Bat09 respective gene packaging signals as described in detail (**Fig. 1A**) [33]. To generate the second MLV with IL-18 expression (Bat09:mH3mN2-NS1-128-IL-18, MLV2), the recombinant porcine IL-18 (rpIL-18) was incorporated between the truncated bat NS1 (NS1-128) and NEP proteins as described previously [39]. Briefly, rpIL-18 was fused to the C-terminal of NS1-128 via a GSGG linker followed by a GSG linker, 2A autoproteolytic cleavage site and NEP (Fig. 1B). NS1-128-IL-18 and NEP are expressed as a single polyprotein, and 2A autoproteolytic cleavage site allows NEP to be released from NS1-128-IL-18 protein during translation. Wild type TX98 H3N2 virus (cluster I) was used as WIV vaccine antigen by UV-inactivation and emulsion in 15% commercial oil-in-water adjuvant at 64 HA unites/dose (Emulsigen D, MVP Laboratories, Inc.). Recently isolated cluster IV H3N2 virus, A/swine/Kansas/10-91088/2010 (KS-91088), was used as challenge virus for *in vivo* pig study [40]. All viruses used as vaccine and challenge strains were propagated in Madin-Darby canine kidney (MDCK) cells.

**FIG 1.**
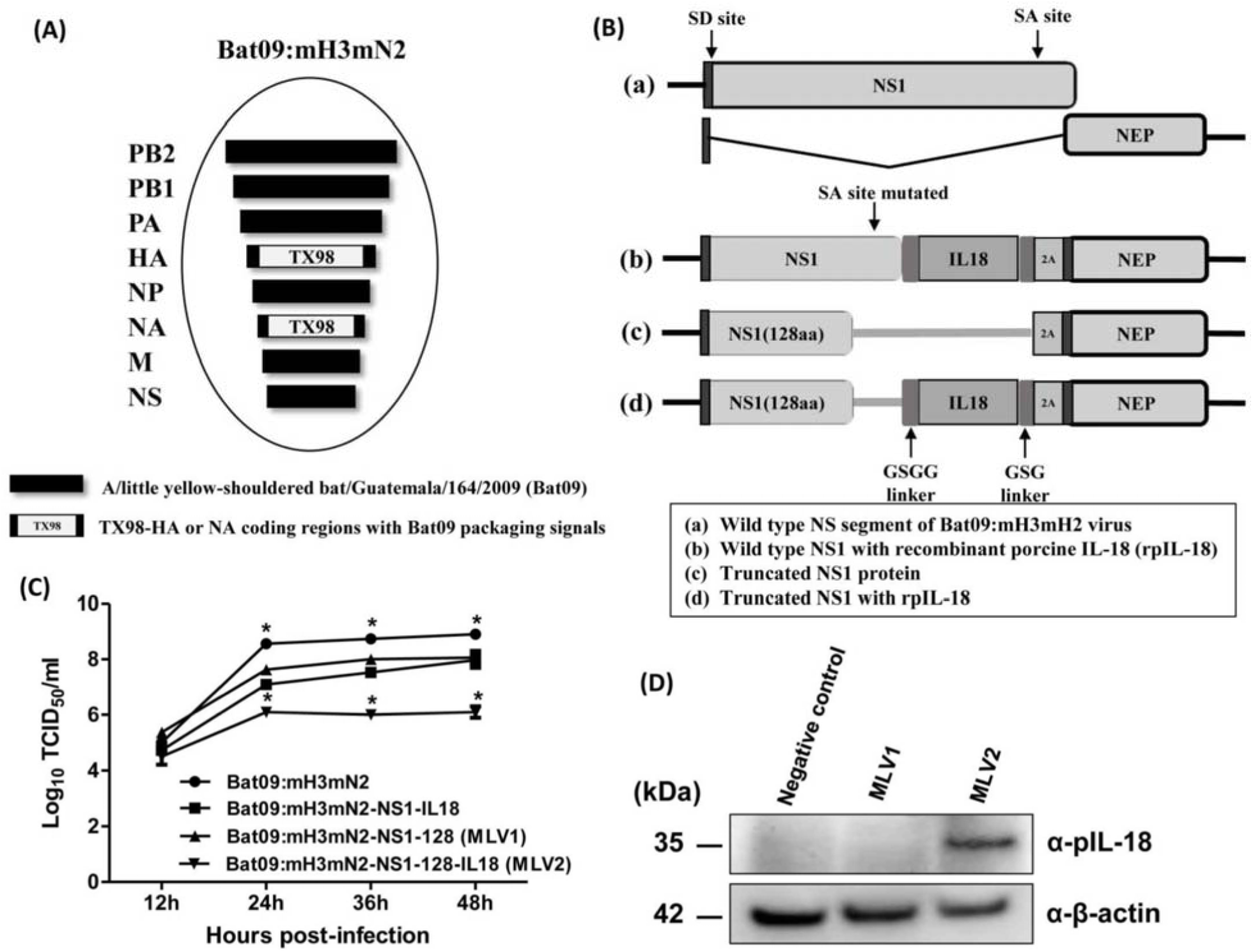
Generation and characterization of MLVs. (A) Chimeric bat influenza virus (Bat09:mH3mH2) which contains six internal genes from the H17N10 A/little yellow-shouldered bat/Guatemala/164/2009 (Bat09) (black bars) and has coding regions of HA and NA from TX98 H3N2 (A/swine/Texas/4199-2/1998, TX98) with Bat09 packaging signals (white bars) was used as replicative vaccine vector to generate MLVs. (B) Wild type NS segment of Bat09:mH3mH2 virus was modified to generate attenuated MLVs. Truncated NS1 protein with 128 amino acids is expressed as a single polyprotein with 2A autoproteolytic cleavage site and NEP. NEP protein is released from NS1 protein during translation. The rpIL-18 was incorporated between NS1 and NEP proteins via GSGG, GSG linkers and 2A autoproteolyitc cleavage site. Splice acceptor site was mutated to inhibit splicing. SD: splice donor site, SA: splice acceptor site. (C) Replication kinetics of recombinant viruses in MDCK cells infected at an MOI of 0.001. Each data point on the curve indicates the means of the results in triplicate, and the error bars indicate standard errors of the mean (SEM). (D) The expression of rpIL-18 with truncated NS1 protein in MLV2 was determined by western blotting analysis from virus infected MDCK cells. Negative control was mock infected control.

### Growth kinetics

To assess the replication kinetics of MLV strains, MDCK cells were cultured in 12-well plates and infected with each virus at a multiplicity of infection (m.o.i.) of 0.01 in triplicate. At 12, 24, 36 and 48h post-infection (hpi), the supernatants of infected cells were collected and virus titers were determined by calculating the 50% tissue culture infective dose (TCID_50_/ml) in MDCK cells using the Reed and Muench method [41].

### Western blotting

Confluent MDCK cells were infected with MLV1 and MLV2 at an m.o.i. of 0.2. Negative control was mock-infected with the PBS. At 24 hpi, infected cells were collected and cell lysates were extracted using CelLytic M cell lysis (Sigma-Aldrich) and extracted cell lysates were loaded to 4-12% Bis-Tris polyacrylamide gel (Invitrogen). The loaded gel was transferred onto a polyvinylidene difluoride (PVDF) membrane (Millipore) and the membrane was blocked using 5% skim milk. The membrane was incubated with a primary mouse anti-pig IL-18 antibody (diluted 1:500, Bio-Rad) or anti-β-actin antibody (diluted 1:500, Santa Cruz) overnight at 4ºC. Horseradish peroxidase (HRP)-conjugated polyclonal rabbit anti-mouse immunoglobulins (diluted 1:1000, room temperature, 1h reaction, Dako) were used as the secondary antibody.

Target protein was detected using SuperSignal West Femto Maximum Sensitivity Substrate according to the manufacturer’s instruction (Thermo Scientific).

### Experimental design

Twenty-three 3-to 4-week-old pigs, which were confirmed to be seronegative for porcine reproductive and respiratory syndrome virus and IAVs by hemagglutination inhibition (HI) assay against currently circulating H3N2 and H1N1 viruses, were used in this study. Pigs were randomly distributed into 4 groups (MLV1, MLV2, WIV vaccinated groups and non-vaccinated control group) and each group contained 6 pigs, while MLV2 vaccinated group had 5 pigs (**Table 1**). The MLV groups were vaccinated with 1.5 ml of 10^6^ TCID_50_/pig by intranasal route once. One group of 6 pigs were administered with 2 ml of 64 HA units of adjuvanted WIV vaccine by intramuscular route and boosted 21 days later with the same dose of the vaccine by the same route. Six pigs in non-vaccinated group (NV) were served as sham vaccinated controls. Pigs were monitored daily for clinical signs and body temperature was measured daily for 7 days post primary vaccination. All pigs were challenged at 28 days post-vaccination (dpv) with 10^5^ TCID_50_/ml of KS-91088 virus by intra-tracheal inoculation (**Table 1**). Rectal temperature and clinical sings were monitored daily after vaccination and challenge. Nasal swabs were collected 0, 3, 5, 7 dpv and 0, 1, 3, 5 days post-challenge (dpc) to evaluate virus nasal shedding. At 0, 14, 28 dpv and 3, 5 dpc, blood samples were collected from each pig for serological analysis. Three pigs from each group were necropsied at 3 and 5 dpc, while two pigs from the MLV2 group were necropsied at 5 dpc (**Table 1**). Bronchoalveolar lavage fluid (BALF) samples were collected by flushing each lung with 50 ml of fresh minimal essential medium (MEM). Virus amounts of nasal swab and BALF samples were determined on MDCK cells by calculating TCID_50_/ml using the Reed and Muench method [41]. To investigate the gene reassortment between MLVs and challenge virus in BALF after challenge, MDCK cells were infected with BALF samples and plaque assay was performed to select a single virus. The purified single virus plaques were amplified for further analysis to identify the origin of each gene segment using gene specific reverse transcription polymerase chain reaction (RT-PCR). RNAs were extracted from each amplified single virus using the QIAamp Viral RNA Mini Kit (Qiagen). Each gene segment was synthesized by gene specific primers (primers available upon request) using SuperScript III One-Step RT-PCR System (Invitrogen).

### Pathological examination of tissues

At necropsy, lungs were removed *in toto* and a single experienced veterinarian assessed the percentage of typical IAV infection gross lesions of each lobe (each lung lobe is considered as 100%) as described previously [42, 43]. The mean of gross lung lesions of seven lung lobes was calculated and the average lung lesions of each pig were presented. Tissue samples of right cardiac lung lobe and trachea from each pig were collected and fixed in 10% neutral buffered formalin immediately after necropsy. They were routinely processed for histopathologic examination, and stained with hematoxylin and eosin at Kansas State Veterinary Diagnostic Laboratory. A certified veterinary anatomic pathologist evaluated the microscopic sections for the presence of lesions. The veterinary anatomic pathologist was blinded to the different treatment groups. Lung scoring was assessed based on our former publication [44]. Microscopic lesions were graded for percentage of airway epithelial necrosis and inflammation (0-4 scale), percentage of airway hyperplasia and regeneration (0-3 scale), percentage of peribronchiolar infiltration (0-3 scale), and percentage of interstitial pneumonia (0-4 scale). Similarly, lesions in the trachea were assessed using a 0-4 scale, based on percentage of degeneration and necrosis of epithelium, and degree of inflammation.

### Antibody detection and ELISA assay

To perform hemagglutination inhibition (HI) assays, heat inactivated sera at 56ºC for 30 min were treated with 20% Kaolin (Sigma-Aldrich) and 0.5% turkey red blood cells (RBCs) to get rid of nonspecific hemagglutination inhibitors and agglutinins. The KS-91088 virus was used as antigen to conduct the HI assay by following standard techniques [45]. Enzyme-linked immunosorbent assays (ELISA) were performed to detect total IgG and IgA antibodies in serum and BLAF against whole virus preparation of KS-91088 as previously described with modifications [46]. Heat inactivated sera were diluted in 5% Fraction V bovine serum albumin (BSA) in PBS at 1:1000 for IgG assay and 1:4 for IgA assay. BALF samples were treated in an equal amount of 10mM dithiothreitol (DTT) solution for 1h at 37ºC for mucus dissociation and dilute in 10% BSA/PBS at 1:1 ratio. All diluted serum and BALF samples were incubated at 37ºC for 1h to absorb nonspecific antibody. Plates were coated with 100 HA unit per 50 μl of KS-91088 virus at room temperature overnight and added 50 μl of each sample per well in triplicate. After 1h incubation, 50 μl of horseradish peroxidase-conjugated goat anti-pig IgG (diluted 1:10000, Bio-Rad) or anti-pig IgA (diluted 1:10000, Bio-Rad) were used to detect antibodies. After 1h incubation with detection antibodies, plates were added with 50 μl of 3,3′,5,5′-Tetramethylbenzidine (TMB) Liquid Substrate System (Sigma-Aldrich) for 10 min followed by adding 50 μl of Stop Reagent for TMB Substrate (Sigma-Aldrich). The optical density (OD) was measured at 450 nm wavelength and antibody levels were analyzed by average of each triplicate samples and reported as the mean of OD values of each group.

### IFN-γ ELISPOT assay

To perform ELISPOT assay for detecting IFN-γ secreting cells (IFN-γ SCs), whole blood samples were collected at 5 dpc to isolate peripheral blood mononuclear cells (PBMCs). ELISPOT plates (MSIPS4510, Millipore) were pre-wetted with 15 μl of 35% ethanol for 1min prior to being coated with 50 μl of 10 μg/ml anti-porcine IFN-γ antibody (BD Biosciences) at 4ºC overnight. Plates were washed 5 times with PBS and blocked with 200 μl of RPMI-1640 medium containing 10% fetal bovine serum (FBS) and 1% antibiotic/antimycotic (Invitrogen) at 37ºC for 2h. The blocked wells were seeded with 100 μl of 10^5^ PBMCs and stimulated with 50 μl of 2 x 10^6^ TCID_50_/ml of UV-inactivated KS-91088 virus. Concanavalin A at 10 μg/ml was used as a positive control and uninfected MDCK media was used as a negative control. After 48h incubation, plates were washed and incubated with 50 μl of 0.5 μg/ml biotinylated-porcine IFN-γ antibody (Invitrogen) for 2h at room temperature. After washing, plates were added with 50 μl of HRP-conjugated Streptavidin (diluted 1:2000, Invitrogen) and incubated for 1h at room temperature. Plates were washed with PBS and added with 100 μl of TMB Membrane Peroxidase Substrate System (KPL) to develop dark blue spots on the membrane. Dry plates were scanned and spots were counted using ImmunoSpot (Cellular Technology) machine and ImmunoCapture (Version 6.3.5) and ImmunoSpot (Version 5.0 Profesional DC) software. The average numbers of counted spots for triplicates well of individual sample were used to present the mean numbers of each group.

### Cytokine and chemokine levels in BALF

Cytokine/Chemokine levels in BALF were quantified by MILLIPLEX MAP Porcine Cytokine/Chemokine Magnetic Bead Panel using the Luminex technology according to the manufacturer’s instructions (Millipore). Each sample was analyzed in triplicate and results presented average values of pigs in each treatment group.

### Statistical analysis

All statistical analysis were conducted using analysis of variance (ANOVA) with a Tukey’s multiple comparison test by GraphPad Prism version 5.0 (GraphPad Software) to compare multiple treatment groups. A *P*-value of 0.05 or less was considered statistically significant.

## Results

### Generation and characterization of modified live-attenuated vaccines

To generate novel MLVs by reverse genetics, the chimeric bat influenza virus (Bat09:mH3mN2), which contains coding regions of HA and NA from TX98 H3N2 virus with Bat09 respective gene packaging signals and six internal genes from the Bat09 virus, was used as a replicative bat influenza vector (**Fig. 1A**). Since LAIVs with a truncated NS1 protein have been shown to provide effective protection against SIV infection with attenuated replication feature in pigs [18, 23], we generated a MLV1 based on Bat09:mH3mN2 virus expressing a truncated NS1 protein of 128 amino acids (Bat09:mH3mN2-NS1-128, MLV1) (**Fig. 1B**). In parallel, MLV2 expressing both a truncated NS1 protein and rpIL-18 was generated (Bat09:mH3mN2-NS1-128-IL-18, MLV2); as IL-18 is associated with inducing a strong cell-mediated immunity (**Fig. 1B**). In addition, the recombinant virus expressing full-length NS1 and rpIL-18 was also produced as a control. Virus replication kinetics of all generated recombinant viruses were evaluated *in vitro*. All recombinant viruses propagated efficiently in Madin-Darby canine kidney (MDCK) cells. All three recombinant viruses (Bat09:mH3mN2-NS1-IL18, Bat09:mH3mN2-NS1-128 and Bat09:mH3mN2-NS1-128-IL-18) were attenuated in vitro as a significantly lower titer was detected at 24, 36 and 48 hours post-infection (hpi) compared to the parental Bat09:mH3mN2 virus (**Fig. 1C**).Interestingly, the viral yield of Bat09:mH3mN2-NS1-128-IL-18 was significantly lower than the other two recombinant viruses at 24, 36 and 48hpi (**Fig. 1C**). The porcine IL-18 expression of Bat09:mH3mN2-NS1-128-IL-18 was confirmed by Western blotting using cell lysates from virus infected MDCK cells (**Fig. 1D**). These results indicate that all three recombinant viruses are attenuated compared to the parental Bat09:mH3mN2 virus and can be used as vaccine candidates. Both MLV1 and MLV2 were selected for further vaccine studies.

### No clinical signs and limited virus shedding found in pigs post vaccination with MLVs

Pigs were inoculated with either MLV1 or MLV2 vaccine candidate by the intranasal route with 10^6^ TCID_50_ per pig. No obvious clinical signs were observed in these pigs post vaccination and prior to challenge. We monitored fever of each pig for 7 days post primary vaccination and no fever was detected in all experimental pigs. To determine whether the MLV1 and MLV2 vaccine candidates shed from vaccinated pigs, nasal swab samples were collected at 3, 5 and 7 days post-vaccination (dpv). Low titer virus shedding was detected in a few pigs only at 3 dpv, not at later time points. Virus was found in 2 out of 6 pigs infected with the MLV1 with a titer of 10^1.7^ or 10^2.3^ TCID_50_/ml, while only 1 out of 5 animals infected with the MLV2 candidate shed virus with a titer of 10^1.7^ TCID_50_/ml. These low amounts of virus detected in a few pigs could be the residual of the inoculated virus.

### Serological response following vaccination with MLV vaccine candidates and WIV

Titers of hemagglutination inhibition (HI) antibodies to the heterologous KS-90188 H3N2 virus were evaluated in serum samples from all groups of vaccinated pigs at 14 dpv, 28 dpv, 3 days post-challenge (dpc) and 5 dpc as depicted in **Table 1**. At 14 dpv, both MLVs induced a low HI antibody titer following a single dose of vaccination, while substantial HI antibody titers were detected from the WIV vaccinated group after booster vaccination at 28 dpv (**Fig. 2A**). However, the HI titers in pigs of all vaccinated groups were lower than 1:40 at all tested time points prior to challenge. Although HI titers from all immunized groups increased to higher than 1:100 following challenge, no significant difference in titers was observed between vaccinated groups (**Fig. 2A**). The non-vaccinated control group had no HI titers before challenge, and lower HI antibody titers were detected following challenge compared to the vaccinated groups. We next investigated IgG levels against KS-91088 H3N2 virus in sera at 28 dpv using a whole-virus ELISA assay. WIV vaccinated pigs produced significantly higher IgG serum levels to KS-91088 than those of MLV vaccinated and non-vaccinated pigs (**Fig. 2B**). There was no significant difference in serum IgG levels between the two MLV vaccinated groups and the non-vaccinated control group. Together, these data show that the two MLV vaccine candidates applied as a single dose and the WIV vaccine applied as two doses induced minimal levels of heterologous HI antibody titers (≤ 40) following vaccination, while the WIV vaccine elicited significantly higher levels of IgG in sera compared to the other groups at 28 dpv.

**Fig 2.**
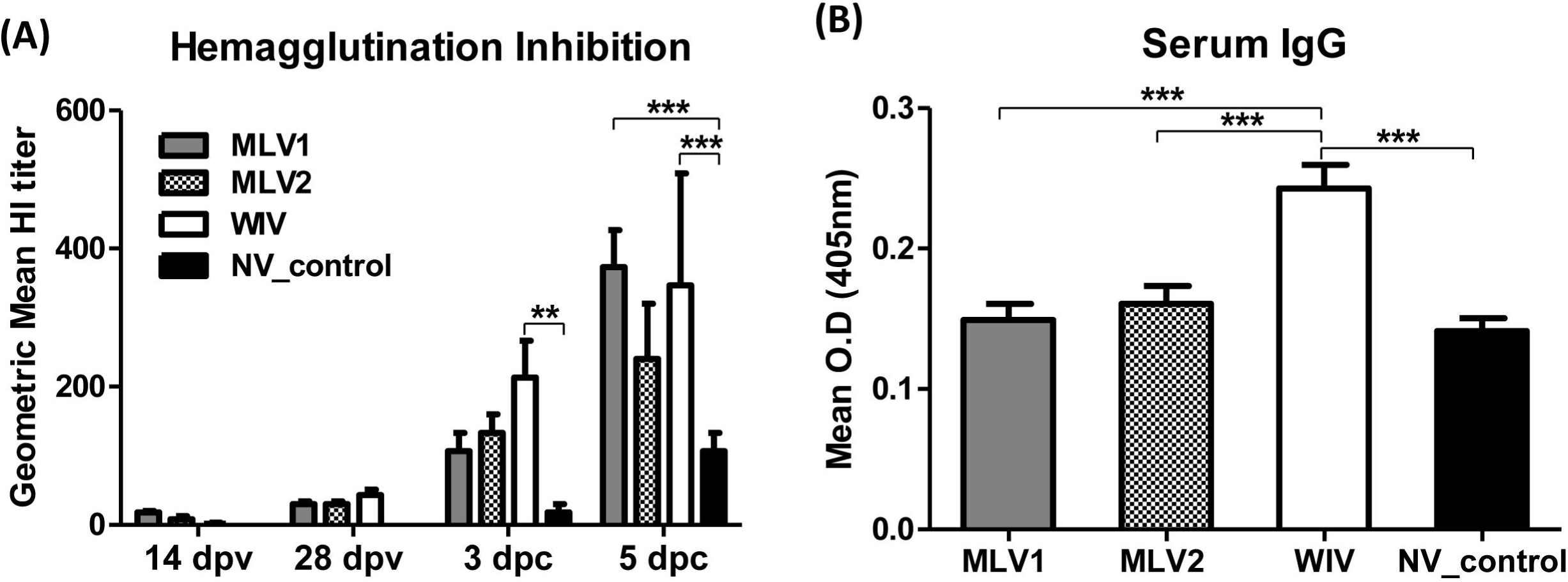
Serum antibody responses in pigs after vaccination. (A) Geometric mean reciprocal titers of hemagglutination inhibition (HI) antibodies to heterologous challenge virus (KS-91088) were determined in sera from all pigs following vaccination and challenge. The reported HI titers are the average titers for each group. (B) At 28 dpv, serum samples were collected from all pigs to evaluate the serum IgG levels against KS-91088 virus using a whole-virus ELISA assay. Antibody levels were analyzed by average of each triplicate sample and reported as the mean of OD values of each group. The error bars indicate standard errors of the mean (SEM). The asterisks (*) represent a statistically significant difference between groups (*: *p*<0.05, **: *p*<0.01 and ***: *p*<0.001).

### Clinical signs, virus replication and shedding, and lung pathology in MLV and WIV vaccinated pigs after challenge

After challenge with heterologous KS-91088 H3N2 virus, all vaccinated pigs as well as the non-vaccinated controls did not show obvious respiratory clinical signs. However, 3 out of 6 pigs immunized with the WIV vaccine showed fever starting at 1 dpc, and all pigs in this group displayed fever at 2 dpc which lasted for 2 days, while no pigs in other groups had fever at 1 dpc. Four or 5 pigs in either the non-vaccinated control group or in the MVL1 group, respectively, showed fever starting at 2 dpc, while all pigs in both groups had fever at 3 dpc. Interestingly, all five pigs in the MLV2 group had fever only at 3 dpc, not at any other days. Pigs immunized with either MLV1 or MLV2 exhibited minimal macroscopic lung lesions with an average of less than 3% (**Fig. 3A and B**). The WIV vaccine group had enhanced macroscopic lung lesions compared to the two MLV groups and the non-vaccination control group at 3 dpc, and the average percentage of lung lesions was significantly higher than the lesions observed with the other groups with an average of more than 50 % (**Fig. 3A and B**). At 5 dpc, no significant differences in lung lesion profiles were observed among the different groups although a higher lung lesion percentage was found in WIV-immunized pigs and minimal lung lesions were found in both MLV vaccinated animals (**Fig. 3B**).

**Fig 3.**
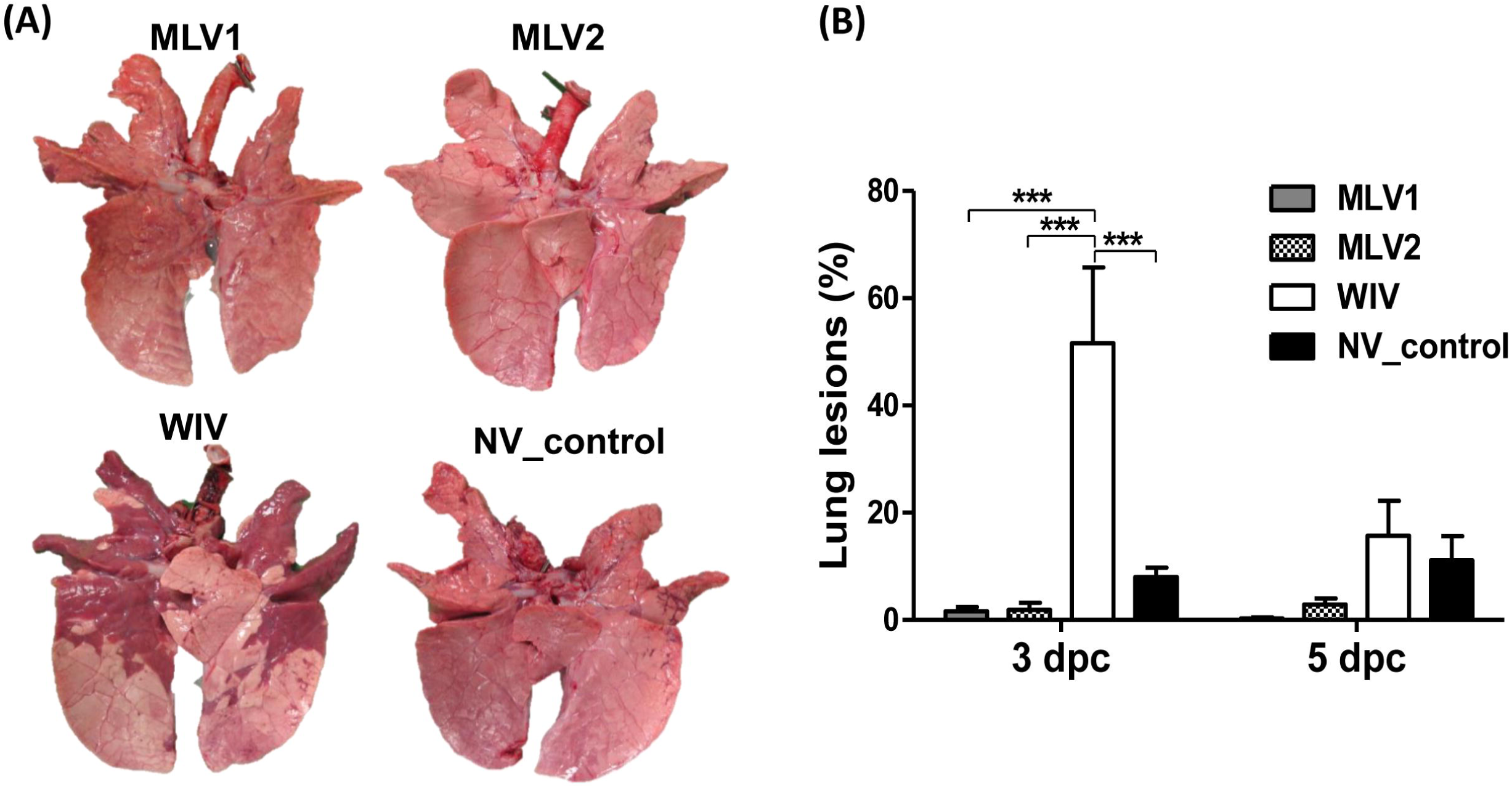
Macroscopic lung lesions in pigs after challenge. (A) Ventral surfaces of lungs from representative pigs in each group at 3 dpc are shown. (B) Macroscopic lung lesions of challenged pigs are presented as the average percentage ± SEM of gross lesions of three pigs in each group at 3 and 5 dpc. The asterisks (*) represent a statistically significant difference between groups (*: *p*<0.05, **: *p*<0.01 and ***: *p*<0.001).

To investigate whether vaccinations could prevent virus replication in lung tissues and limit virus shedding in pigs, we determined virus titers in bronchoalveolar lavage fluid (BALF) and nasal swab samples. At 3 dpc, virus was detected in BALF samples collected from pigs of each group except for one pig from the MLV1 group. In contrast to WIV-vaccinated pigs, a lower virus titer was detected in BALF samples of other three groups of pigs. However, a significant difference in virus titers was only observed between the MLV2-vaccinated and the WIV-vaccinated groups (**Fig. 4A**). At 5 dpc, virus was not detected in BALF from pigs receiving either the MLV1 or MLV2 candidate vaccines, whereas virus was detected inone out of three pigs in the WIV vaccine group and from all 3 pigs in the non-vaccinated control group (**Fig. 4A**). No virus was detected in nasal samples collected from all pigs at 1 dpc (**Fig. 4B**). At 3 dpc, virus was detected in nasal swabs collected from 5 out of 6 pigs in the MLV1 group as well as from all pigs in other 3 groups. A significantly lower titer was found in the MLV1 group compared to those of the other 3 groups at this time point post challenge. No virus was detected in nasal swabs of 2 pigs immunized with the MLV2 vaccine candidate at 5 dpc, whereas virus was found in all pigs in either the WIV or non-vaccinated control groups (**Fig. 4B**). In contrast, significant less virus was detected in 2 out of 3 nasal swabs collected in the MVL1 immunized pigs at 5 dpc. In summary, both MLV vaccine candidates reduced virus replication in lungs and nasal shedding after challenge with a heterologous H3N2 virus; this is in contrast to both the WIV vaccinated and non-vaccinated groups (**Fig. 4**).

**Fig 4.**
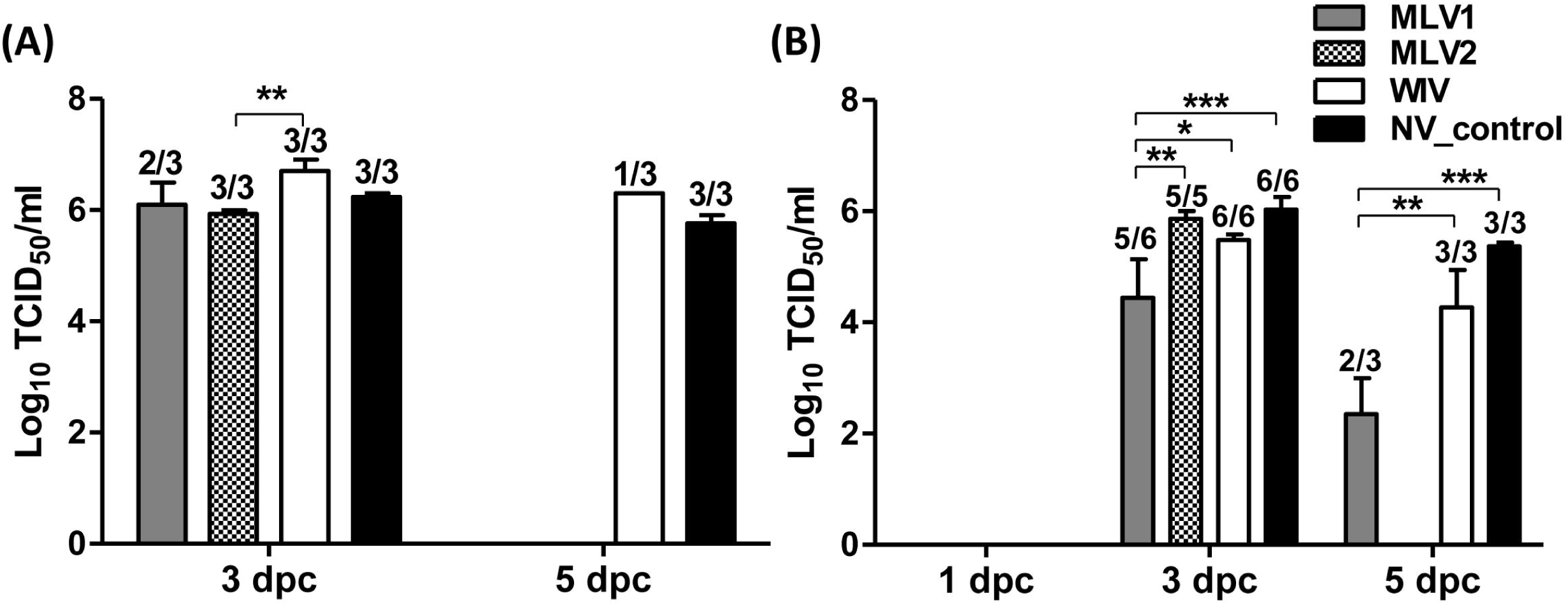
Virus replication in BALF and nasal shedding in nasal swabs after challenge. Mean of virus titers in BALF (A) and in nasal swabs (B) from challenged pigs were evaluated on the days indicated. Virus titers were determined by calculating the 50% tissue culture infective dose (TCID_50_)/ml in MDCK cells. The number of pigs with positive virus isolation out of the total number of tested pigs is presented above of each bar. The asterisks (*) represent a statistically significant difference between groups (*: *p*<0.05, **: *p*<0.01 and ***: *p*<0.001).

Histopathological lesions were examined in lungs and trachea collected from pigs at 5 dpc. Microscopic lung lesion scores were not significantly different between the groups, however, a lower score trend was observed in pigs immunized with either the MLV1 or MLV2 vaccine candidates (**Fig. S1 A**). The MLV2 group and the non-vaccinated control group demonstrated the statistically less microscopic damages to the trachea when compared to the WIV vaccine group (**Fig. S1 B**). However, there was no significant difference in microscopic trachea lesions between the two MLV groups and the non-vaccinated control groups. Similar to the above described enhanced macroscopic lung lesions, the WIV vaccine group exhibited more severe histopathological lung and trachea damages as characterized by marked peribrochiolar lymphocytic cuffing, degenerated airway epithelium, extensive infiltration of inflammatory cells within the lumen of airways, attenuated mucosa of trachea and infiltration of lymphocytes and plasma cells in the lamina propria of the trachea with transepethelial migration of inflammatory cells (**Fig. S1 C**). These data suggest that both MLV vaccine candidates can prevent virus replication in lungs and limit virus transmission more efficiently than the WIV vaccine when challenged with heterologous virus in the absence of enhanced lung pathology and disease (i.e., VAERD).

### Assessment of reassortment between MLVs and challenge virus in BALF

Previous studies have shown that chimeric bat influenza viruses fail to reassort with conventional IAVs by co-infection experiments [33–35]. Herein, we investigated reassortment events between MLVs and challenge virus in BALF samples post challenge. Sixty and ninety single virus plaques isolated from BALF of the MLV1 group and MLV2 group, respectively, were purified and the origin of the gene segments of each isolated virus was determined using gene specific RT-PCR assays. The results showed that all segments of the 150 tested isolates belong to the KS-91088 challenge virus and no reassortant viruses with the MLV candidate vaccines were detected.

### Antigen-specific T-cell response after challenge

To evaluate cell-mediated immune response after challenge, peripheral blood mononuclear cells (PBMCs) were isolated from blood collected from pigs at 5 dpc. An ELISPOT assay was performed to detect antigen-specific IFN-γ secreting cells (IFN-γ SCs) in response to heterologous KS-91088 antigens. All immunized pigs showed significantly higher numbers of antigen-specific IFN-γ SCs compared to the non-vaccinated control pigs (**Fig. 5A**). However, both MLV candidate vaccines induced significantly greater antigen-specific IFN-γ recall responses to the heterologous KS-91088 antigens when compared to the WIV vaccinated pigs (**Fig. 5A**).

**Fig 5.**
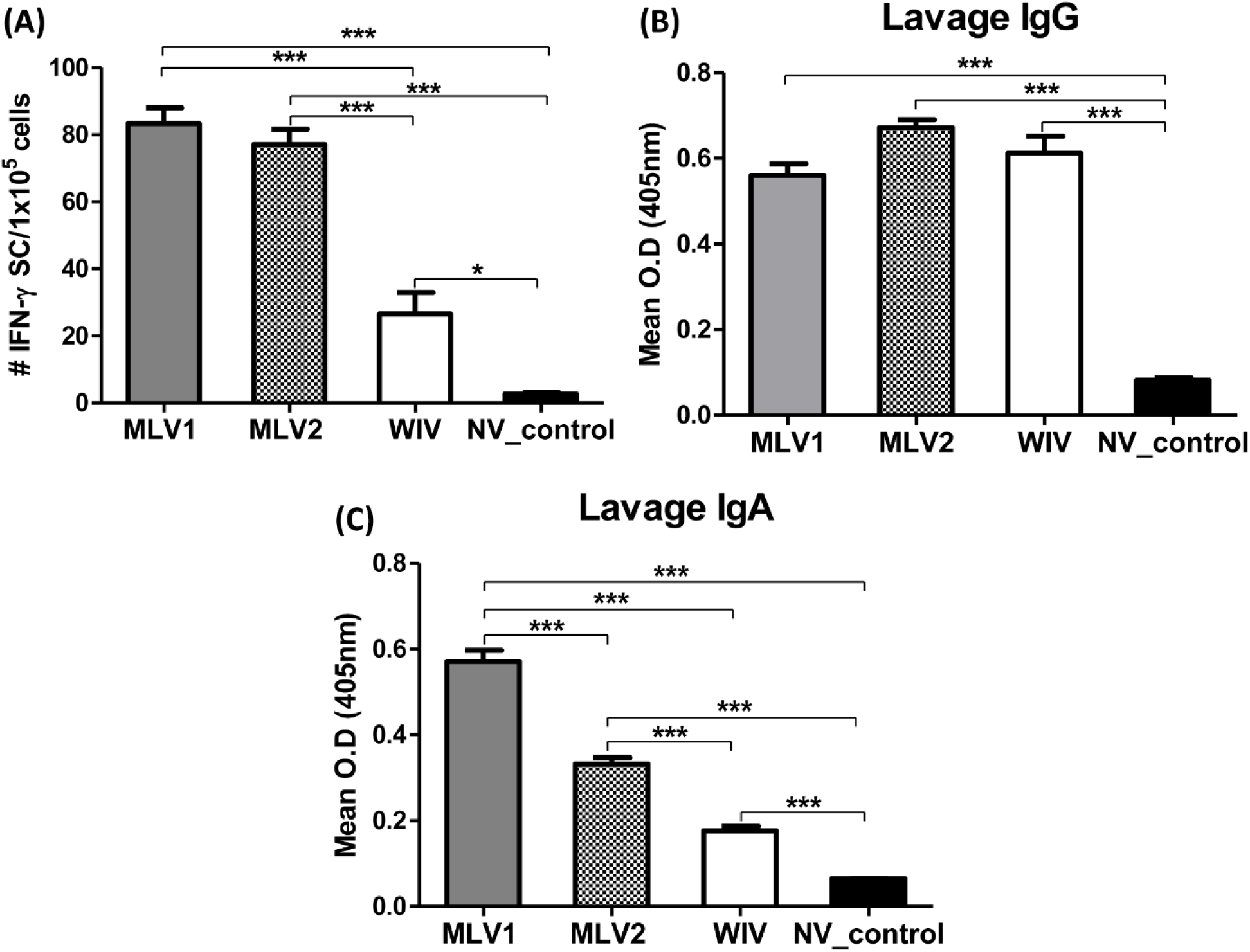
Antigen-specific T-cell response in PBMCs and antibody response to heterologous challenge virus in BLAF at 5 dpc. (A) PBMCs were collected from pigs at 5 dpc and ELISPOT assay was performed to detect antigen-specific IFN-γ secreting cells (IFN-γ SCs) which respond to heterologous KS-91088 antigen. The average numbers of counted spots for triplicates well of individual sample were used to present the data of the mean numbers of each group. Levels of KS-91088 specific IgG (B) and IgA (C) antibodies in BALF at 5 dpc were evaluated by ELISA. Antibody levels were analyzed by average of each triplicate sample and expressed as the mean of OD values of each group. The error bars indicate standard errors of the mean (SEM). The asterisks (*) represent a statistically significant difference between groups (*: *p*<0.05, **: *p*<0.01 and ***: *p*<0.001).

### Antibody response to challenge virus in BALF samples

The levels of IgG and IgA antibodies reactive to the KS-91088 challenge virus in BALF at 5 dpc were measured using an isotype-specific ELISA. The results showed that there were significantly higher levels of IgG in BALF from all vaccinated pigs compared to non-vaccinated control pigs (**Fig. 5B**). However, both MLVs and the WIV vaccine elicited similar levels of cross-reactive IgG antibodies in BALF samples. Similar to the IgG levels in BALF, the KS-91088 reactive IgA levels of all vaccinated pigs were significantly higher than those of non-vaccinated pigs (**Fig. 5C**). In contrast, significantly higher levels of IgA antibodies were detected in BALF from pigs immunized with either MLV1 or MLV2 compared to the WIV vaccine group (**Fig. 5C**).

### Cytokine and chemokine levels in BALF following challenge

Pulmonary levels of cytokines and chemokines in BALF collected from pigs at 3 and 5 dpc were assessed using the Luminex technology. All measured cytokine and chemokine levels in BALF from the WIV vaccine group were significantly higher than those detected from both the MLV1 and MLV2 groups as well as the non-vaccinated control group at either 3 or 5 dpc **(Fig. S2)**. Significant differences in all tested cytokine and chemokine levels between the MLV groups and the non-vaccinated control group were not observed except for IL-6 and IL-12 at 3 dpc (**Fig. S2**).

## Discussion

In North America, WIV vaccines are widely used in swine herds to prevent and control swine influenza. However, the complexity and diverse ecology of SIVs lead to repeated failure of WIV vaccines even if farm-specific autogenous WIV vaccines are used [7, 47]. Furthermore, even though WIV vaccines are able to provide adequate protection against homologous influenza virus infection, ineffective protection together with induction of VAERD are observed when WIV vaccinated pigs are infected with antigenically mismatched strains [8–11]. Therefore, this limited efficacy of traditional WIV vaccines is not adequate to provide broad protection against co-circulating antigenically rather diverse SIVs. LAIVs using a variety of genetic modifications have been shown to provide superior protection against homologous and heterologous SIV challenges compared to WIV vaccines, without causing VAERD [17, 25, 46, 48]. However, there are safety concerns of using LAIVs that potentially reassort with circulating SIV strains. In the present study, we developed new MLV vaccine candidates for swine influenza using an attenuated chimeric bat influenza virus as a vaccine vector and assessed their efficacy against heterologous virus challenge in swine. The novel MLV vaccines conferred better protection than the WIV vaccine against heterologous virus challenge as evidenced by reduced virus replication in lungs and limiting nasal virus shedding (**Fig. 4**). Although significantly reduced virus replication and shedding was observed in the MLV vaccinated pigs at 5 dpc, virus was still present at 3 dpc **(Fig. 4)**. The lack of early protection against respiratory virus replication in the MLV groups may be due to the high, intra-tracheal challenge dose (10^5^TCID_50_/pig) used in this study and/or the antigenic mismatch between challenge and vaccine strains. Even if the average macroscopic and microscopic lung lesions were not significantly different between the MLV groups and the non-vaccinated control group, reduced lung damage was observed in pigs immunized with MLVs compared to the non-vaccinated control group (**Fig. S1**). Most importantly, VAERD was only observed in WIV vaccinated pigs but not in pigs immunized with the MLV candidate vaccines upon heterologous challenge **(Fig. 3 and Fig. S1)**. Moreover, no reassortment was detected between MLVs and the challenge virus in BALF after challenge. These data indicate that the bat influenza vectored MLV vaccine candidates can provide protection from heterologous SIV infection without safety concerns compared to traditional WIV and LAIV vaccines by overcoming the risks of VAERD and virus reassortment.

VAERD is a major adverse effect of WIV vaccines when immunized pigs are infected with heterologous virus, especially HA mismatched virus. The WIV vaccine used in this study was derived from TX98 (H3N2) virus which is a triple-reassortant cluster I of H3 virus, whereas the HA of the KS-91088 (H3N2) challenge virus belongs to the cluster IV of H3 virus that has no or limited cross-reactivity with the vaccine strain. The identity of the HA protein sequences between the TX98 and KS-91088 viruses is approximately 91% and this high divergence of HA between the WIV vaccine and the challenge KS-91088 virus most likely resulted in VAERD with severe and exacerbated lung damages **(Fig. 3 and Fig. S1)**. Previous studies demonstrated that VAERD correlated with the presence of high level of non-neutralizing antibodies and the absence of neutralizing antibodies against antigenically mismatched virus [12, 14, 46, 49]. In particular, it has been reported that cross-reactive HA2 antibodies which target the HA2 domain but not the HA1 globular head domain contributed to enhanced IAV infection in cells through promoting the fusion process between virus and cell membrane, and this infection enhancement is associated with VAERD [46, 49]. In the present study, all vaccine groups displayed similar low levels of heterologous HI antibody titers (≤ 40) before challenge, even after the WIV vaccine group was boosted with a second dose **(Fig. 2A)**. However, high cross-reactive IgG levels against the KS-91088 virus were detected in sera from WIV vaccinated pigs while the MLV vaccinated groups produced minimal levels of serum IgG antibodies similar to those of the non-vaccinated group at 28 dpv **(Fig. 2B)**. Previous studies reported similar results such as limited HI response to heterologous virus strains in vaccinated groups prior to challenge, and more cross-reactive serum IgG antibodies elicited in WIV vaccine groups when compared to LAIV groups [17, 46, 50]. Moreover, previous studies found that sera from WIV vaccinated pigs contained high titers of cross-reactive antibodies against the HA2 domain of a heterologous challenge virus; this was consistent with high cross-reactive serum IgG levels and low titers of cross-reactive HA1 antibodies which corresponded with limited HI antibody response and low serum neutralizing antibodies to a heterologous challenge virus [46]. This suggests that the high cross-reactive IgG serum antibodies present in WIV vaccinated pigs prior to challenge observed in this study could reflect the high level of fusion-enhancing HA2 antibodies against the heterologous challenge virus; in addition, the low levels of heterologous HI titers may correspond to the low level of virus-neutralizing antibodies. If this is correct, the high anti-HA2 antibodies in WIV vaccinated pigs might enhance virus replication in epithelial cells and this might result in longer duration as well as higher titers of lung virus replication and nasal virus shedding in the WIV vaccine group when compared to the MLV groups **(Fig. 4)**. The enhanced lung virus replication may cause the influx of numerous inflammatory immune cells followed by excessive inflammatory reactions with high pulmonary cytokine/chemokine levels **(Fig. S2)**. This scenario could be the basis for the severe lung damage and the development of VAERD in WIV vaccinated animals; in contrast, the MLV groups do not develop elevated fusion-enhancing HA2 antibodies and therefore, did not develop VAERD when challenged with a heterologous virus **(Fig. 3 and Fig. S1)**.

Several reports have shown that intranasally administered LAIVs are able to elicit cross-reactive T-cell responses and robust mucosal antibodies which play an important role in LAIV efficacy for heterologous challenge protection [9, 23, 25, 48, 51, 52]. In this study, MLV vaccine candidates induced significantly more antigen-specific IFN-γ SCs in the blood in response to heterologous antigen as well as significantly higher levels of cross-reactive IgA antibodies in BALF compared to WIV vaccinated pigs after challenge; interestingly post-challenge heterologous HI titers in sera and IgG levels in BALF were similar among all vaccinated groups **(Fig. 2A and Fig. 5)**. This suggests that significant levels of cross-reactive cell-mediated and mucosal immune responses which were induced by the MLV vaccines might contribute to providing better cross-protection for heterologous challenge when compared to a WIV vaccine. Cell-mediated immunity is mediated by T-cell responses including CD4^+^ T-cells and CD8^+^ T-cells. Since the majority of T-cell responses are directed towards epitopes which are highly conserved between different influenza virus strains, T-cell immunity has the potential to elicit heterologous cross-protective immune responses [53–55]. In pigs, it has been reported that T-cell mediated immunity conferred cross-protection to heterologous challenge with reduced disease severity [56, 57]. However, bat influenza viruses have a distant genetic relation to classical IAVs with sharing only50-70% genetic identity [29, 30]. Therefore, the degree of conservation of viral epitopes between bat influenza viruses and conventional IAVs and whether viral proteins of bat influenza viruses are able to induce cell-mediated immunity as well as cross-protection to classical IAV infection remains to be investigated. The present study demonstrated that the novel MLV vaccine candidates were able to elicit robust cross-reactive IgA antibodies in BALF even with a single intranasal vaccination **(Fig. 5C)**. Since IgG antibody levels in BALF were similar between all vaccine groups, mucosal cross-reactive IgA antibodies were most likely crucial for the local immunity in lungs to limit virus replication and reduce lung pathology under heterologous challenge conditions **(Fig. 3 and Fig. 4A)**. This is an agreement with other studies which showed an important role of local IgA immunity induced by intranasally administered LAIVs for SIV challenge infections [9, 25, 46, 48]. In a previous study, Masic *et al*. described that a two dose LAIV vaccination was able to induce increased antigen-specific serum HI, IgG and mucosal IgA antibodies compared to a one dose of intranasal LAIV vaccination [48]. Therefore, further studies will be needed to investigate whether a boosting regimen of bat influenza vectored MLV vaccine candidates can induce improved immune responses and a better protection from subsequent SIV infection in pigs when compared to a single dose regimen.

IL-18 which is secreted from macrophages, neutrophils, dendritic cells, T cells and epithelial cells and has been shown to up-regulate IFN-γ expression by activating natural killer (NK) cells as well as T lymphocytes [36, 38]. Previous studies have demonstrated that influenza virus infection can induce IL-18 expression in human alveolar macrophages and IL-18 expression was necessary for optimal cytokine production and adequate protection against influenza infections [37, 38, 58-60]. In addition, one recent study reported that mucosal associated invariant T cells (MAITs) were activated and produced antiviral effectors including IFN-γ via a IL-18 dependent mechanism during influenza infection and contributed to antiviral influenza immunity [61]. However, IL-18 expression in young piglets up to one month old is significantly reduced at their respiratory mucosal epithelium, which is the major infection site of IAVs [62]. Therefore, we hypothesized that exogenous IL-18 expression in conjunction with vaccine administration may help to provide enhanced protective immunity by inducing strong CMI response to SIV infection and we investigated the immunomodulatory effects of rpIL-18 which was incorporated into the MLV2 vaccine. However, no significant difference was observed in vaccine efficacy between two MLV vaccine candidates, regardless of rpIL-18 expression. We assume that the MLV2 virus (Bat09:mH3mN2-NS1-128-IL-18) did not replicate as efficiently as needed to express enough IL-18 to show an obvious functional effect of IL-18 **(Fig. 1C and 1D)**. Further studies will be required to investigate the effect of IL-18 on vaccine efficacy using less attenuated MLVs, a higher dose of vaccine or a booster regimen.

In this present study, we developed two new MLV candidate vaccines using attenuated chimeric bat influenza virus as a vaccine vector and clearly demonstrated their protective immunity against SIV infection. One advantage of bat influenza vectored MLV vaccines is the mode of intranasal immunization which mimics natural infection. It induces not only protective local immunity including IgA production at the site of natural IAV infection but also a broad cross-protective CMI without causing VAERD. The major concern of LAIV vaccination is the risk of gene reassortment with circulating strains and the generation of novel IAVs. However, chimeric bat influenza viruses have not been able to reassort with conventional IAVs by co-infection experiments and no reassortment of genomic segments between the MLVs used here and the challenge virus was observed in BALF [33–35]. This indicates that bat influenza vectored MLV vaccines can be used as safe and efficacious vaccine vectors without the potential risk of gene reassortment with conventional IAVs. Additionally, some internal viral proteins of bat influenza viruses have been shown to be functionally compatible with conventional IAV proteins by reverse genetics [33-35, 63]. Thus, we anticipate that some conserved viral epitopes of bat influenza viruses in the MLV vaccines could induce some degree of cross-reactive immunity; other vectored vaccines can express only a limited number of viral proteins mainly related to inducing neutralizing antibodies, such as HA or NA [25]. Taken together, bat influenza vectored MLV candidate vaccines provided protective immunity against heterologous challenge and can be used as safe and efficacious live virus vaccines to prevent swine influenza infections in pigs and might be also used in other species.

## Conclusions

Novel MLV vaccines using an attenuated chimeric bat influenza virus were able to reduce virus replication and pathology in lungs, and nasal shedding following a heterologous virus challenge and induce robust mucosal and T-cell immune responses in pigs without VAERD. These results demonstrate that bat influenza vectored MLV vaccines are effective and safe to protect swine from SIV infection without risks of VAERD and reassortment.

## Acknowledgments

The authors thank staffs from the Comparative Medicine Group at Kansas State University for supporting the animal studies, and Jennifer Phinney from the Kansas State Veterinary Diagnostic Lab for technical assistance with H&E and IHC staining. This work was supported by grants from NIH NIAID 1R01AI134768-01A1 and the NIAID-funded Center of Excellence for Influenza Research and Surveillance (CEIRS, #HHSN272201400006C). The funders had no role in study design, data collection and analysis, decision to publish, or preparation of the manuscript.

## Author contribution

WM conceived and designed the experiments. JL, YL, YL, AGC, MD, YL, JM, SS, JAR and WM performed the experiments. JL, DB, AGC, JAR and WM analyzed the data. JL and WM wrote the paper with input from all other authors.

## Supplemental Figure legends

**Fig S1. Microscopic lesions of lung and trachea in pigs at 5 dpc**. Microscopic scores of lung (A) and trachea (B) are presented as mean scores ± SEM of pig in each group at 5 dpc. The asterisks (*) represent a statistically significant difference between groups (*: *p*<0.05). (C) Lung and trachea sections of pigs at 5 dpc were stained with H&E. Lungs from pigs in MLVs groups are moderately affected and contain few infiltrates of inflammatory cells within the lumen of airways and mild to moderate lymphocytic and plasmacytic peribronchiolar cuffing (red asterisks). Lungs from pigs in WIV and NV-control groups are severely affected with infiltrates of inflammatory cells intermixed with fibrin and edema fluid within the lumen of airways and marked peribronchiolar cuffing of lymphocytes, plasma cells, and neutrophils (red asterisks) that are migrating through the epithelium. The airway epithelium is markedly attenuated and degenerated (arrows). Trachea from MLV2 immunized pig is mildly affected with little perivascular infiltrates of lymphocytes and plasma cells. The mucosa is normal. Tracheal mucosa of pigs in MLV1 and NV-control groups are moderately hyperplastic (M) and there are moderate infiltrates of lymphocytes and plasma cells in the lamina propria (red asterisk). In WIV group, the mucosa of the trachea is severely attenuated (arrow) and there is infiltration of lymphocytes and plasma cells in the lamina propria which extends deep to the glands (red arrow) with transepithelial migration of inflammatory cells. Bars = 20 um.

**Fig S2. Cytokine and chemokine levels in BALF after challenge**. The expression levels of porcine cytokine/chemokines in BALF following challenge were quantified using the Luminex technology. Data represent the average values ± SEM of pigs in each group on the days indicated. The asterisks (*) indicate a statistically significant difference between virus infected groups (*: *p*<0.05. **: *p*<0.01 and ***: *p*<0.001).

## References

[1] Abente EJ, Santos J, Lewis NS, Gauger PC, Stratton J, Skepner E, et al. The Molecular Determinants of Antibody Recognition and Antigenic Drift in the H3 Hemagglutinin of Swine Influenza A Virus. J Virol 2016 Sep 15;90(18):8266–80.

[2] Vincent A, Awada L, Brown I, Chen H, Claes F, Dauphin G, et al. Review of influenza A virus in swine worldwide: a call for increased surveillance and research. Zoonoses Public Health 2014 Feb;61(1):4–17.

[3] Nelson MI, Stratton J, Killian ML, Janas-Martindale A, Vincent AL. Continual Reintroduction of Human Pandemic H1N1 Influenza A Viruses into Swine in the United States, 2009 to 2014. J Virol 2015 Jun;89(12):6218–26.

[4] Rajao DS, Gauger PC, Anderson TK, Lewis NS, Abente EJ, Killian ML, et al. Novel Reassortant Human-Like H3N2 and H3N1 Influenza A Viruses Detected in Pigs Are Virulent and Antigenically Distinct from Swine Viruses Endemic to the United States. J Virol 2015 Nov;89(22):11213–22.

[5] Lindstrom S, Garten R, Balish A, Shu B, Emery S, Berman L, et al. Human infections with novel reassortant influenza A(H3N2)v viruses, United States, 2011. Emerg Infect Dis 2012 May;18(5):834–7.

[6] Smith GJ, Vijaykrishna D, Bahl J, Lycett SJ, Worobey M, Pybus OG, et al. Origins and evolutionary genomics of the 2009 swine-origin H1N1 influenza A epidemic. Nature 2009 Jun 25;459(7250):1122–5.

[7] Van Reeth K, Ma W. Swine influenza virus vaccines: to change or not to change-that’s the question. Curr Top Microbiol Immunol 2013;370:173–200.

[8] Vincent AL, Ciacci-Zanella JR, Lorusso A, Gauger PC, Zanella EL, Kehrli ME, Jr., et al. Efficacy of inactivated swine influenza virus vaccines against the 2009 A/H1N1 influenza virus in pigs. Vaccine 2010 Mar 24;28(15):2782–7.

[9] Loving CL, Lager KM, Vincent AL, Brockmeier SL, Gauger PC, Anderson TK, et al. Efficacy in pigs of inactivated and live attenuated influenza virus vaccines against infection and transmission of an emerging H3N2 similar to the 2011-2012 H3N2v. J Virol 2013 Sep;87(17):9895–903.

[10] Vincent AL, Lager KM, Janke BH, Gramer MR, Richt JA. Failure of protection and enhanced pneumonia with a US H1N2 swine influenza virus in pigs vaccinated with an inactivated classical swine H1N1 vaccine. Vet Microbiol 2008 Jan 25;126(4):310–23.

[11] Reeth KV, Brown I, Essen S, Pensaert M. Genetic relationships, serological cross-reaction and cross-protection between H1N2 and other influenza A virus subtypes endemic in European pigs. Virus Res 2004 Jul;103(1-2):115–24.

[12] Gauger PC, Vincent AL, Loving CL, Lager KM, Janke BH, Kehrli ME Jr., et al. Enhanced pneumonia and disease in pigs vaccinated with an inactivated human-like (delta-cluster) H1N2 vaccine and challenged with pandemic 2009 H1N1 influenza virus. Vaccine 2011 Mar 24;29(15):2712–9.

[13] Gauger PC, Vincent AL, Loving CL, Henningson JN, Lager KM, Janke BH, et al. Kinetics of lung lesion development and pro-inflammatory cytokine response in pigs with vaccine-associated enhanced respiratory disease induced by challenge with pandemic (2009) A/H1N1 influenza virus. Vet Pathol 2012 Nov;49(6):900–12.

[14] Rajao DS, Chen H, Perez DR, Sandbulte MR, Gauger PC, Loving CL, et al. Vaccine-associated enhanced respiratory disease is influenced by haemagglutinin and neuraminidase in whole inactivated influenza virus vaccines. J Gen Virol 2016 Jul;97(7):1489–99.

[15] Souza CK, Rajao DS, Loving CL, Gauger PC, Perez DR, Vincent AL. Age at Vaccination and Timing of Infection Do Not Alter Vaccine-Associated Enhanced Respiratory Disease in Influenza A Virus-Infected Pigs. Clin Vaccine Immunol 2016 Jun;23(6):470–82.

[16] Loving CL, Vincent AL, Pena L, Perez DR. Heightened adaptive immune responses following vaccination with a temperature-sensitive, live-attenuated influenza virus compared to adjuvanted, whole-inactivated virus in pigs. Vaccine 2012 Aug 31;30(40):5830–8.

[17] Vincent AL, Ma W, Lager KM, Richt JA, Janke BH, Sandbulte MR, et al. Live attenuated influenza vaccine provides superior protection from heterologous infection in pigs with maternal antibodies without inducing vaccine-associated enhanced respiratory disease. J Virol 2012 Oct;86(19):10597–605.

[18] Solorzano A, Webby RJ, Lager KM, Janke BH, Garcia-Sastre A, Richt JA. Mutations in the NS1 protein of swine influenza virus impair anti-interferon activity and confer attenuation in pigs. J Virol 2005 Jun;79(12):7535–43.

[19] Masic A, Babiuk LA, Zhou Y. Reverse genetics-generated elastase-dependent swine influenza viruses are attenuated in pigs. J Gen Virol 2009 Feb;90(Pt 2):375–85.

[20] Pena L, Vincent AL, Ye J, Ciacci-Zanella JR, Angel M, Lorusso A, et al. Modifications in the polymerase genes of a swine-like triple-reassortant influenza virus to generate live attenuated vaccines against 2009 pandemic H1N1 viruses.J Virol 2011 Jan;85(1):456–69.

[21] Coleman JR, Papamichail D, Skiena S, Futcher B, Wimmer E, Mueller S. Virus attenuation by genome-scale changes in codon pair bias. Science 2008 Jun 27;320(5884):1784–7.

[22] Kaplan BS, Souza CK, Gauger PC, Stauft CB, Robert Coleman J, Mueller S, et al. Vaccination of pigs with a codon-pair bias de-optimized live attenuated influenza vaccine protects from homologous challenge. Vaccine 2018 Feb 14;36(8):1101–7.

[23] Vincent AL, Ma W, Lager KM, Janke BH, Webby RJ, Garcia-Sastre A, et al. Efficacy of intranasal administration of a truncated NS1 modified live influenza virus vaccine in swine. Vaccine 2007 Nov 19;25(47):7999–8009.

[24] Tatsis N, Fitzgerald JC, Reyes-Sandoval A, Harris-McCoy KC, Hensley SE, Zhou D, et al. Adenoviral vectors persist in vivo and maintain activated CD8+ T cells: implications for their use as vaccines. Blood 2007 Sep 15;110(6):1916–23.

[25] Braucher DR, Henningson JN, Loving CL, Vincent AL, Kim E, Steitz J, et al. Intranasal vaccination with replication-defective adenovirus type 5 encoding influenza virus hemagglutinin elicits protective immunity to homologous challenge and partial protection to heterologous challenge in pigs. Clin Vaccine Immunol 2012 Nov;19(11):1722–9.

[26] Wesley RD, Tang M, Lager KM. Protection of weaned pigs by vaccination with human adenovirus 5 recombinant viruses expressing the hemagglutinin and the nucleoprotein of H3N2 swine influenza virus. Vaccine 2004 Sep 3;22(25-26):3427–34.

[27] Vander Veen RL, Loynachan AT, Mogler MA, Russell BJ, Harris DL, Kamrud KI. Safety, immunogenicity, and efficacy of an alphavirus replicon-based swine influenza virus hemagglutinin vaccine. Vaccine 2012 Mar 2;30(11):1944–50.

[28] Vander Veen RL, Mogler MA, Russell BJ, Loynachan AT, Harris DL, Kamrud KI. Haemagglutinin and nucleoprotein replicon particle vaccination of swine protects against the pandemic H1N1 2009 virus. Vet Rec 2013 Oct 12;173(14):344.

[29] Tong S, Li Y, Rivailler P, Conrardy C, Castillo DA, Chen LM, et al. A distinct lineage of influenza A virus from bats. Proc Natl Acad Sci U S A 2012 Mar 13;109(11):4269–74.

[30] Tong S, Zhu X, Li Y, Shi M, Zhang J, Bourgeois M, et al. New world bats harbor diverse influenza A viruses. PLoS Pathog 2013;9(10):e1003657.

[31] Li Q, Sun X, Li Z, Liu Y, Vavricka CJ, Qi J, et al. Structural and functional characterization of neuraminidase-like molecule N10 derived from bat influenza A virus. Proc Natl Acad Sci U S A 2012 Nov 13;109(46):18897–902.

[32] Sun X, Shi Y, Lu X, He J, Gao F, Yan J, et al. Bat-derived influenza hemagglutinin H17 does not bind canonical avian or human receptors and most likely uses a unique entry mechanism. Cell Rep 2013 Mar 28;3(3):769–78.

[33] Zhou B, Ma J, Liu Q, Bawa B, Wang W, Shabman RS, et al. Characterization of uncultivable bat influenza virus using a replicative synthetic virus. PLoS Pathog 2014 Oct;10(10):e1004420.

[34] Juozapaitis M, Aguiar Moreira E, Mena I, Giese S, Riegger D, Pohlmann A, et al. An infectious bat-derived chimeric influenza virus harbouring the entry machinery of an influenza A virus. Nat Commun 2014 Jul 23;5:4448.

[35] Yang J, Lee J, Ma J, Lang Y, Nietfeld J, Li Y, et al. Pathogenicity of modified bat influenza virus with different M genes and its reassortment potential with swine influenza A virus. J Gen Virol 2017 Apr;98(4):577–84.

[36] Dinarello CA. IL-18: A TH1-inducing, proinflammatory cytokine and new member of the IL-1 family. J Allergy Clin Immunol 1999 Jan;103(1 Pt 1):11–24.

[37] Denton AE, Doherty PC, Turner SJ, La Gruta NL. IL-18, but not IL-12, is required for optimal cytokine production by influenza virus-specific CD8+ T cells. Eur J Immunol 2007 Feb;37(2):368–75.

[38] Takeda K, Tsutsui H, Yoshimoto T, Adachi O, Yoshida N, Kishimoto T, et al. Defective NK cell activity and Th1 response in IL-18-deficient mice. Immunity 1998 Mar;8(3):383–90.

[39] Manicassamy B, Manicassamy S, Belicha-Villanueva A, Pisanelli G, Pulendran B, Garcia-Sastre A. Analysis of in vivo dynamics of influenza virus infection in mice using a GFP reporter virus. Proc Natl Acad Sci U S A 2010 Jun 22;107(25):11531–6.

[40] Ma J, Shen H, Liu Q, Bawa B, Qi W, Duff M, et al. Pathogenicity and transmissibility of novel reassortant H3N2 influenza viruses with 2009 pandemic H1N1 genes in pigs. J Virol 2015 Mar;89(5):2831–41.

[41] Reed LJ, Muench H. A simple method of estimating fifty per cent endopoints. American Journal of Epidemiology 1938 1 MAY 1938;27(3):493–7.

[42] Richt JA, Lager KM, Janke BH, Woods RD, Webster RG, Webby RJ. Pathogenic and antigenic properties of phylogenetically distinct reassortant H3N2 swine influenza viruses cocirculating in the United States. Journal of clinical microbiology 2003 Jul;41(7):3198–205.

[43] Ma W, Vincent AL, Gramer MR, Brockwell CB, Lager KM, Janke BH, et al. Identification of H2N3 influenza A viruses from swine in the United States. Proceedings of the National Academy of Sciences of the United States of America 2007 Dec 26;104(52):20949–54.

[44] Lee J, Henningson J, Ma J, Duff M, Lang Y, Li Y, et al. Effects of PB1-F2 on the pathogenicity of H1N1 swine influenza virus in mice and pigs. J Gen Virol 2017 Jan;98(1):31–42.

[45] WHO. WHO manual on animal influenza diagnosis and surveilance. Geneva, Switzerland: WHO, 2002.

[46] Gauger PC, Loving CL, Khurana S, Lorusso A, Perez DR, Kehrli ME, Jr., et al. Live attenuated influenza A virus vaccine protects against A(H1N1)pdm09 heterologous challenge without vaccine associated enhanced respiratory disease. Virology 2014 Dec;471-473:93–104.

[47] Vincent AL, Ma W, Lager KM, Janke BH, Richt JA. Swine influenza viruses a North American perspective. Adv Virus Res 2008;72:127–54.

[48] Masic A, Lu X, Li J, Mutwiri GK, Babiuk LA, Brown EG, et al. Immunogenicity and protective efficacy of an elastase-dependent live attenuated swine influenza virus vaccine administered intranasally in pigs. Vaccine 2010 Oct 8;28(43):7098–108.

[49] Khurana S, Loving CL, Manischewitz J, King LR, Gauger PC, Henningson J, et al. Vaccine-induced anti-HA2 antibodies promote virus fusion and enhance influenza virus respiratory disease. Sci Transl Med 2013 Aug 28;5(200):200ra114.

[50] Sandbulte MR, Platt R, Roth JA, Henningson JN, Gibson KA, Rajao DS, et al. Divergent immune responses and disease outcomes in piglets immunized with inactivated and attenuated H3N2 swine influenza vaccines in the presence of maternally-derived antibodies. Virology 2014 Sep;464-465:45–54.

[51] Cheng X, Zengel JR, Suguitan AL, Jr., Xu Q, Wang W, Lin J, et al. Evaluation of the humoral and cellular immune responses elicited by the live attenuated and inactivated influenza vaccines and their roles in heterologous protection in ferrets. J Infect Dis 2013 Aug 15;208(4):594–602.

[52] Sun K, Ye J, Perez DR, Metzger DW. Seasonal FluMist vaccination induces cross-reactive T cell immunity against H1N1 (2009) influenza and secondary bacterial infections. J Immunol 2011 Jan 15;186(2):987–93.

[53] Soema PC, van Riet E, Kersten G, Amorij JP. Development of cross-protective influenza a vaccines based on cellular responses. Front Immunol 2015;6:237.

[54] Sridhar S. Heterosubtypic T-Cell Immunity to Influenza in Humans: Challenges for Universal T-Cell Influenza Vaccines. Frontiers in immunology 2016;7:195.

[55] Kees U, Krammer PH. Most influenza A virus-specific memory cytotoxic T lymphocytes react with antigenic epitopes associated with internal virus determinants. J Exp Med 1984 Feb 1;159(2):365–77.

[56] Kappes MA, Sandbulte MR, Platt R, Wang C, Lager KM, Henningson JN, et al. Vaccination with NS1-truncated H3N2 swine influenza virus primes T cells and confers cross-protection against an H1N1 heterosubtypic challenge in pigs. Vaccine 2012 Jan 5;30(2):280–8.

[57] Van Reeth K, Braeckmans D, Cox E, Van Borm S, van den Berg T, Goddeeris B, et al. Prior infection with an H1N1 swine influenza virus partially protects pigs against a low pathogenic H5N1 avian influenza virus. Vaccine 2009 Oct 23;27(45):6330–9.

[58] Liu B, Mori I, Hossain MJ, Dong L, Takeda K, Kimura Y. Interleukin-18 improves the early defence system against influenza virus infection by augmenting natural killer cell-mediated cytotoxicity. J Gen Virol 2004 Feb;85(Pt 2):423–8.

[59] Pirhonen J, Sareneva T, Kurimoto M, Julkunen I, Matikainen S. Virus infection activates IL-1 beta and IL-18 production in human macrophages by a caspase-1-dependent pathway. J Immunol 1999 Jun 15;162(12):7322–9.

[60] Pirhonen J, Sareneva T, Julkunen I, Matikainen S. Virus infection induces proteolytic processing of IL-18 in human macrophages via caspase-1 and caspase-3 activation. Eur J Immunol 2001 Mar;31(3):726–33.

[61] Loh L, Wang Z, Sant S, Koutsakos M, Jegaskanda S, Corbett AJ, et al. Human mucosal-associated invariant T cells contribute to antiviral influenza immunity via IL-18-dependent activation. Proc Natl Acad Sci U S A 2016 Sep 6;113(36):10133–8.

[62] Muneta Y, Goji N, Tsuji NM, Mikami O, Shimoji Y, Nakajima Y, et al. Expression of interleukin-18 by porcine airway and intestinal epithelium. J Interferon Cytokine Res 2002 Aug;22(8):883–9.

[63] Aydillo T, Ayllon J, Pavlisin A, Martinez-Romero C, Tripathi S, Mena I, et al. Specific Mutations in the PB2 Protein of Influenza A Virus Compensate for the Lack of Efficient Interferon Antagonism of the NS1 Protein of Bat Influenza A-Like Viruses. J Virol 2018 Apr 1;92(7).

