## Supplemental materials for "Bat influenza vectored NS1-truncated live vaccine protects pigs against heterologous virus challenge"

**Running title: Bat influenza vectored live vaccine protects pigs**

### Jinhwa Lee^a^, Yonghai Li^a^, Yuhao Li^a^, A. Giselle Cino-Ozuna^a^, Michael Duff^a^, Yuekun Lang^a^, Jingjiao Ma^a^, Sunyoung Sunwoo^a^, Juergen A. Richt^a^, Wenjun Ma^a,b,c*^

**Supplemental Figures**

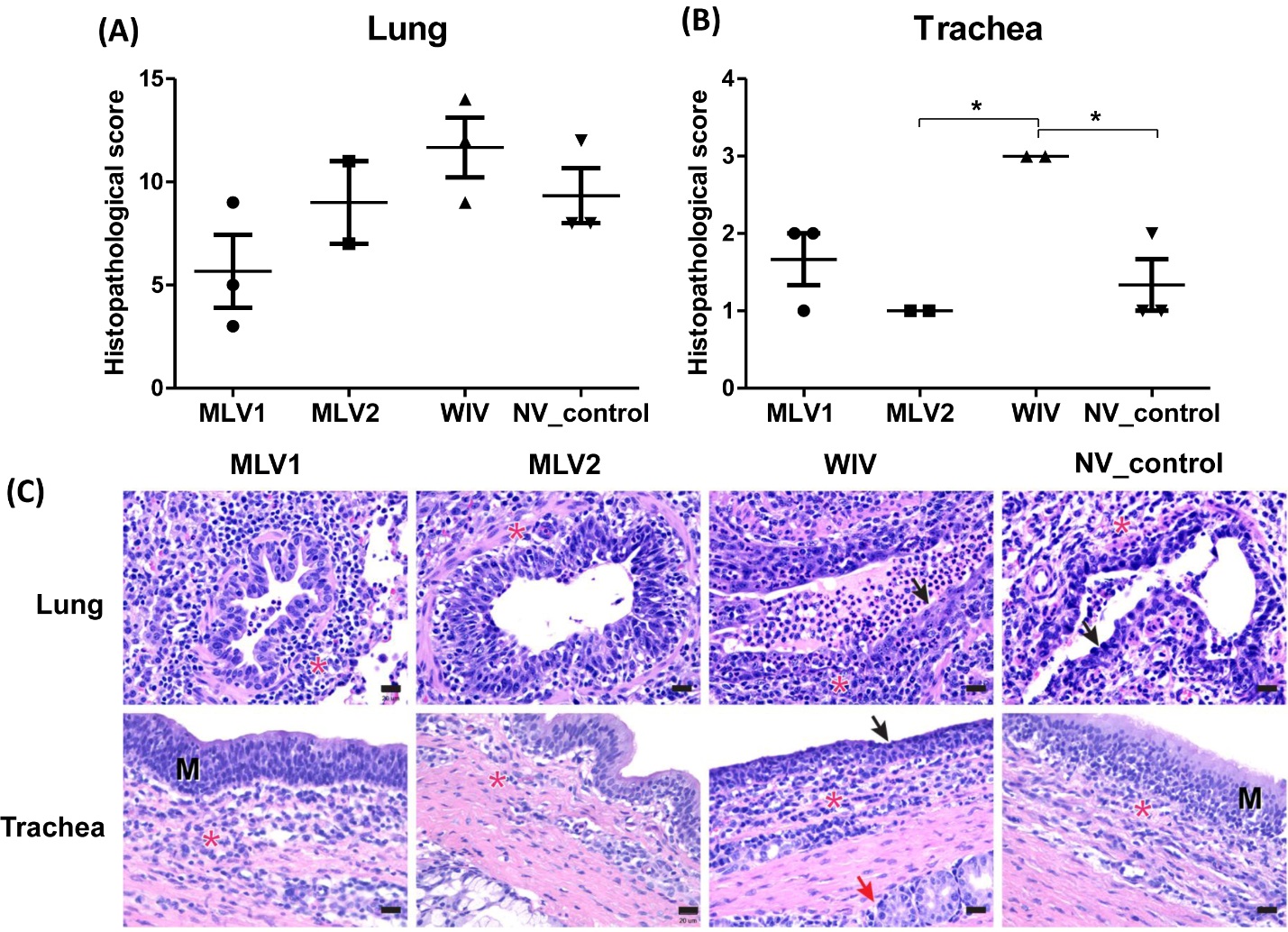

**Fig S1. Microscopic lesions of lung and trachea in pigs at 5 dpc.** Microscopic scores of lung (A) and trachea (B) are presented as mean scores ± SEM of pig in each group at 5 dpc. The asterisks (*) represent a statistically significant difference between groups (*: *p*<0.05). (C) Lung and trachea sections of pigs at 5 dpc were stained with H&E. Lungs from pigs in MLVs groups are moderately affected and contain few infiltrates of inflammatory cells within the lumen of airways and mild to moderate lymphocytic and plasmacytic peribronchiolar cuffing (red asterisks). Lungs from pigs in WIV and NV-control groups are severely affected with infiltrates of inflammatory cells intermixed with fibrin and edema fluid within the lumen of airways and marked peribronchiolar cuffing of lymphocytes, plasma cells, and neutrophils (red asterisks) that are migrating through the epithelium. The airway epithelium is markedly attenuated and degenerated (arrows). Trachea from MLV2 immunized pig is mildly affected with little perivascular infiltrates of lymphocytes and plasma cells. The mucosa is normal. Tracheal mucosa of pigs in MLV1 and NV-control groups are moderately hyperplastic (M) and there are moderate infiltrates of lymphocytes and plasma cells in the lamina propria (red asterisk). In WIV group, the mucosa of the trachea is severely attenuated (arrow) and there is infiltration of lymphocytes and plasma cells in the lamina propria which extends deep to the glands (red arrow) with transepithelial migration of inflammatory cells. Bars = 20 um.

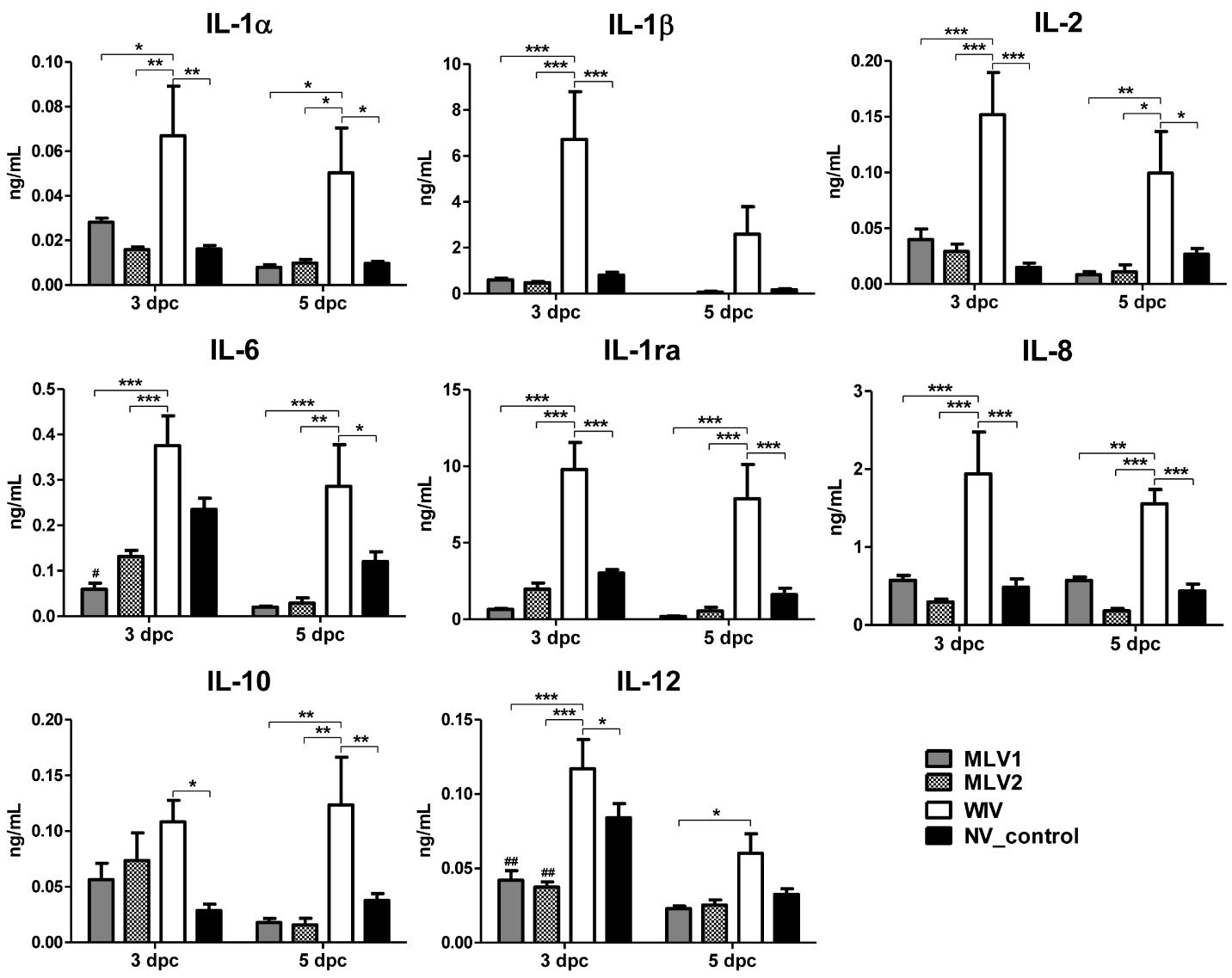

**Fig S2.** **Cytokine and chemokine levels in BALF after challenge.** The expression levels of porcine cytokine/chemokines in BALF following challenge were quantified using the Luminex technology. Data represent the average values ± SEM of pigs in each group on the days indicated. The asterisks (*) indicate a statistically significant difference between virus infected groups (*: *p*<0.05. **: *p*<0.01 and ***: *p*<0.001).
